# The ENDS of assumptions; an online tool for the Epistemic Nonparametric Drug-response Scoring

**DOI:** 10.1101/2021.12.01.470849

**Authors:** Ali Amiryousefi, Bernardo Williams, Mohieddin Jafari, Jing Tang

## Abstract

**Motivation:** The drugs sensitivity analysis is often elucidated from drug dose-response curves. These curves capture the degree of cell viability (or inhibition) over a range of induced drugs, often with parametric assumptions that are rarely validated.

**Results:** We present a class of nonparametric models for the curve fitting and scoring of drug dose-responses. To allow a more objective representation of the drug sensitivity, these epistemic models devoid of any parametric assumptions attached to the linear fit, allow the parallel indexing such as IC_50_ and AUC. Specifically, three nonparametric models including Spline, Monotonic, and Bayesian (npS, npM, npB) and the parametric Logistic (pL) are implemented. Other indices including Maximum Effective Dose (MED) and Drug-response Span Gradient (DSG) pertinent to the npS are also provided to facilitate the interpretation of the fit. The collection of these models are implemented in an online app, standing as useful resource for drug dose-response curve fitting and analysis.

**Availability:** The ENDS is freely available online at https://irscope.shinyapps.io/ENDS/ and source codes can be obtained from https://github.com/AmiryousefiLab/ENDS.

**Supplementary information:** Supplementary data are available at *Bioinformatics* and https://irscope.shinyapps.io/ENDS/

**Contributions:** AA conceived the study and developed the models, AA and BW adopted and implemented the methods, JT provided the funding, AA, BW, MJ, and JT wrote the paper.

## 1 Introduction

Most statistical methodology for dose-response studies has been using the parametric models such as Log-logistic, Gaussian, Weibull or Gompertz functions (Holland-Letz and Kopp-Schneider, 2015). The most frequently used method has been logistic fitting (Berkson, 1944), particularly the four parameter logistic model by weighted least squares (pL) (Vølund, 1978). pL fit albeit powerful, relies on its assumptions regarding the data such as normality of residuals while the continuity of the explanatory variable is often unmet or challenging in under-sampled studies pertinent to the drug-responses (Montgomery *et al*., 2021). These not only limit the application of these models but also affect the derived indices such as half-maximal inhibitory concentration (IC_50_) and Area Under Curve (AUC) as important measures in determination of the drug response, for example in drug synergy testing (Tang *et al*., 2015). Therefore, the application of other measures according to statistical notions such as unimodality and homoscedasticity have been also proposed to evaluate drug sensitivity (Jafari *et al*., 2021). Despite the common use of the pL in the drug response scoring studies with a number of R packages devoted for its implementation, e.g. the drc package (Ritz *et al*., 2015), yet the use of those for less advanced users can pose complications. To provide a more objective characterization of drug sensitivity, we propose an online collection of epistemic nonparametric drug-response scoring (ENDS) models and indices. This class of models bereft of assumptions regarding the model fitting and its parameters, is appealing for its simplicity and intuitiveness and hence allows more objective characterization from the experiment. In turn, these models pivot more on the experiment results than compromising the accuracy in case of deviation from assumptions underlying the parametric models, expanding more room for more intrinsic data exploration.

## 2 Materials and methods

The ENDS web app is fully coded in R and is freely accessible at https://irscope.shinyapps.io/ENDS/. Following the Github link provided above, the source code is available for more versatile use by advanced users. The web app also provides the theoretical accounts of the implemented models and related indices (Sup 1). A single-cell level colorectal cancer drug dose response dataset is used for illustrating the functionality of the tool (Roerink *et al*., 2018).

### 2.1 Models, data input, and usage

The ENDS encompasses four models: **1) nonparametric Spline (npS)**, which connects the mean or median of each dose-response. This simple model is accompanied with novel indices including Drug Span Gradient (DSG) and Min-Max Viability Band (MMB) that facilitate more characterization from the fit (Sup 1.1.1, 1.2.3, Fig. 1A). **2) nonparametric Monotonic (npM)**, which is an isotonic regression fit of the npS with non-decreasing condition (or non-increasing in case of percentage inhibition, (Sup 1.1.2; Fig. 1B). **3) nonparametric Bayesian (npB)**, which is an heuristic model fit by a normal Cumulative Distribution Function with a choice of a prior distribution (Sup 1.1.3; Fig. 1B), and **4) parametric Logistic (pL)**, which is a conventional four parameter sigmoid fit (Sup 1.1.4; Fig. 1B). The input of the ENDS is a single .*csv* file including doses as rows and samples as columns (Sup 1.2.1). The ENDS by default will generate the npS. Users can add npM, npB, and pL model fits and choose the display of more indices for example with outliers exclusion option off or a spectrum choice of IC values (Sup 1.2.3).

**Fig. 1.**
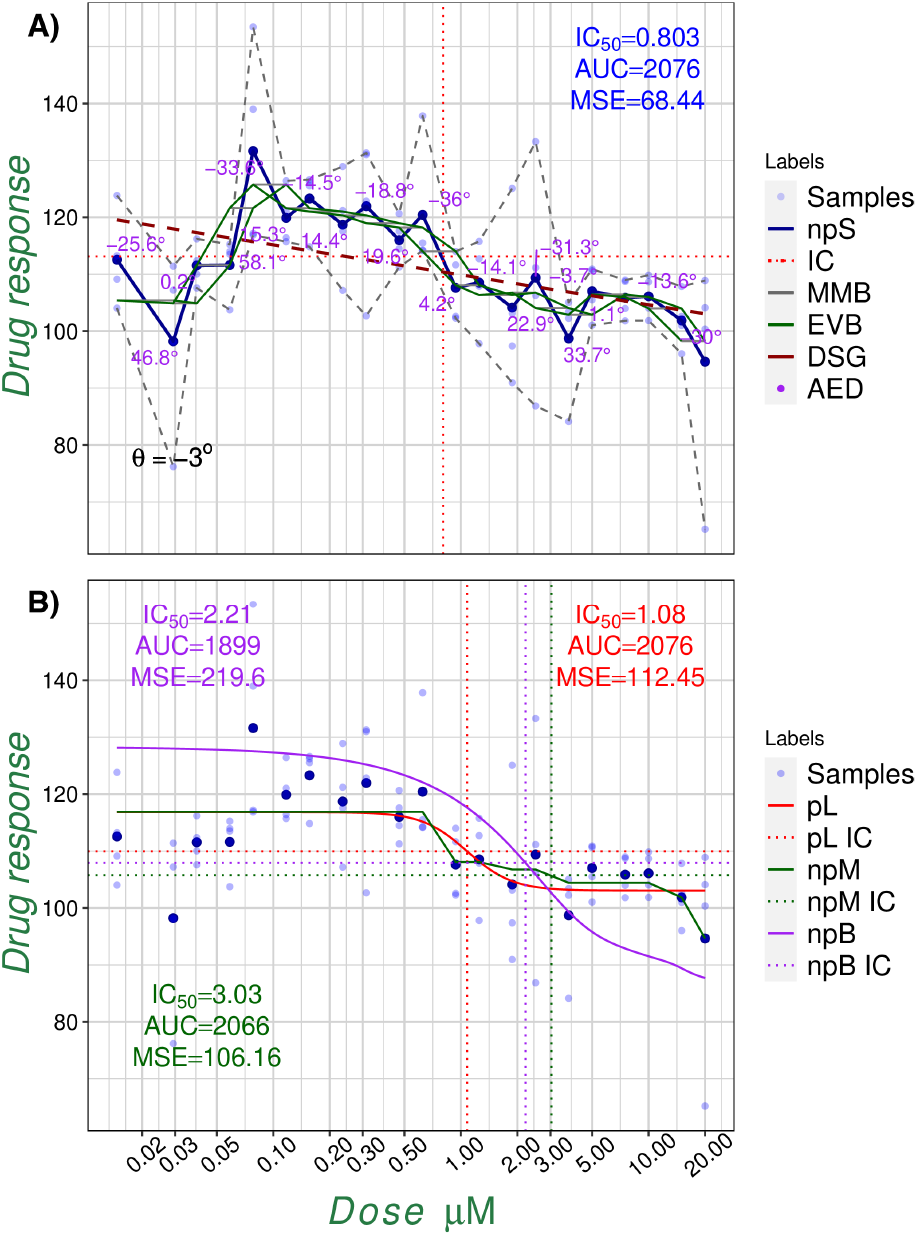
Examples of the ENDS model fitting. The overall outputs of the ENDS models overlaid with the respective IC_50_, AUC, and MSE for the 5-FU drug of the first patient sample of the colorectal cancer study (Roerink et al., 2018). Due to different mathematical formulas for each fit, each model is providing a unique IC_50_, AUC and MSE values. A) The npS model on the mean of the samples, Absolute Efficiency Degrees (AEDs), Min-Max and Empirical Viability Bands (MMB, EVB), next to the Drug Span Gradient (DSG, *θ* = −3°) and Maximum Effective Dose (MED=0.85*μM* with −33.6° gradient). B) The npM, npB, and pL fits for the same data.

## 3 Results

Emphasizing on the nonparametric paradigm, the ENDS provides an online platform for drug dose-response scoring and model fitting. Three fundamentally important yet intrinsically different nonparametric models are introduced that along with other optional scores such as choice of median for each sample, provide a useful tool for drug dose-response analysis. The ENDS also provides the resultant high resolution downloadable plots in different formats (Sup 1.2.2, 1.2.3; Fig. 1).

## 4 Discussion and conclusion

The ENDS presents mathematically justified models and and have a rigorous theory that supports their use. Despite their nonparametric nature, each model is exhibiting various degrees of simplicity and are intrinsically different. At one end, the npS is intuitive with no internal assumption, while the more heuristic npB is rather computationally costly and may be biased by the choice of prior for small datasets (Sup 1.1.1). The ENDS also provides the pL fitting and scores. This allows to compare the performance of the models *a posteriori*. For example, our survey over all the available data of the colorectal study (Roerink *et al*., 2018) revealed the least mean IC_50_ and Mean Square Error (MSE) of the npS fits. While the former indicates the statistically significant over-estimation of the IC_50_ by pL and npB (Sup 2), the latter is indicating the significantly better fit to the data (Sup Tab. 3). Observing no significant difference between the means of the AUC for different models (Sup Tab. 4) next to the least AUC variance for npB, prioritizes this model over the rest, however this is challenged with the lower mean and variance MSE of the other models (Sup Fig. 2). Despite different performance of each model, it is the experimental settings and the level of plausibility of each model that shall guide the users with their preferences. For non-parametric models, we believe that the ENDS can stand as a valuable tool to complement the existing applications.

## Acknowledgements

We thank the European Research Council (ERC) starting grant DrugComb (Informatics approaches for the rational selection of personalized cancer drug combinations, No. 716063) and Academy of Finland grant No. 332454 for the financial support. *Conflict of interest:* none declared.

## 1 Supplementary Material

In this section we will go through the technical details of the ENDS, the functionality of the web application available at https://irscope.shinyapps.io/ENDS/, a detailed technical explanation of the models and a comparative analysis which was obtained by running the models through the collection of drug-response data found in Roerink *et al*. (2018).

### 1.1 Models

This section provides the mathematical account of the four models presented in the paper.

#### 1.1.1 Nonparametric Spline (*npS*)

The model is the collection of linear functions that connect the average responses at each dose. Given a sample of doses *x*_1_, ‥, *x*_*n*_ where *x*_*i*_ ≤ *x*_*i*+1_ and for each i-th dose we have *m* responses *y*_*i*1_, ‥, *y*_*im*_, then for each dose we can obtain the dose 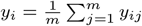 or alternatively we can calculate the dose medians. The simple spline that connects each of the means with a linear function is given by the piece-wise linear function

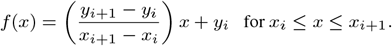

The function is defined on the interval *x*_1_ ≤ *x* ≤ *x*_*n*_. Note that this function is not monotonic necessarily, so if we define the IC_50_ as the value in the *x*-axis such that *f* (*IC*_50_) = (max *y* + min *y*)*/*2, then it might not be unique as it crosses the above function on multiple points, if the function is constant on an interval *I* such that *f* (*I*) = (max *y*+min *y*)*/*2, then IC_50_ = inf *I* is the infimum of the values on the interval. Thus, we calculate all the possible *x*-axis values that map to the halfway point of the responses and select as the effective dose 50% the one closest in absolute value to the one found by the monotonic fit. For any value *p* ∈ (0, 1) the IC_*p*_ will be the chosen out of the candidates *f* (*IC*_*p*_) = min *y* + *p*(max *y* −min *y*), such that it is closest in distance to the uniquely value obtained by the monotone fit. Note that, if the function is constant we again choose as the IC_*p*_ the infinum of the interval.

In order to obtain the angles associated to the slope at each dose we have the *i*th angle as *θ*_*i*_ = *arctan*((*y*_*i*+1_ − *y*_*i*_)*/*(*x*_*i*+1_ − *x*_*i*_)). Note that if the npS is calculated on the means at each dose then the squared error 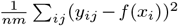 will be minimized since the mean minimizes the L2 norm, alternatively if we calculate npS using the medians then the L1 norm would be minimized.

#### 1.1.2 Nonparametric Monotone (*npM*)

This nonparametric model is fit over the mean responses at each dose or the medians and it is analogous to npS, with the difference of forcing a non-increasing constrain on the connected linear functions. If the spline between two doses does not have a non-positive slope, then the average of the previous doses is recursively calculated until the next spline has a non-positive slope, which connects previously calculated average with the next data point.

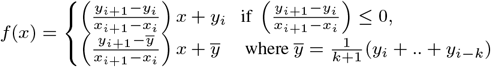

where *k* is the smallest integer such that the slope is non-positive, this is 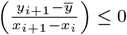. It has been shown that this fit is the one that minimizes the squared errors ∥*y* −*f* (*x*)∥^2^ with the constrain that *f* (*x*_*i*_) ≥ *f* (*x*_*i*+1_). This fit will produce a non increasing fit, even in cases where the pL model or the npB produce an increasing fit.

For the calculation of the *p*-th percent max inhibitory concentration *p* ∈ (0, 1) we find the unique value in the *x*-axis such that

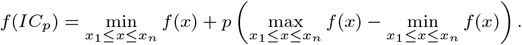

#### 1.1.3 Nonparametric Bayesian (*npB*)

This nonparametric model connects a mixture of normal CDFs with Bayesian modeling to solve for the posterior parameters. The parameters knots *K* and variance *λ* are given *a priori. K* regulates where the distribution functions will be centered and *λ* the variance of the distributions, we choose *K* to be the doses available and *lambda* is found by grid search, chosen as the one with least square error. Namely, given the points (*x*_*i*_, *y*_*i*_) for *i* = 1‥*n*, with knots 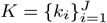 and parameter *λ >* 0 we will model the relation between *x* and *y* with the Bayesian hierarchical model;

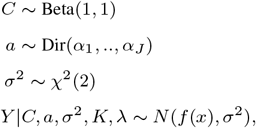

where

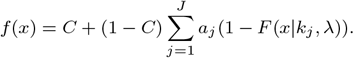

The weights *a* = [*a*_1_, ‥, *a*_*n*_] are non-negative and sum up to one, and *F* is the cumulative distribution function of a Gaussian random variable with mean *k*_*j*_ and variance *λ*. We can use MCMC to sample from the posterior distribution

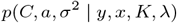

An in-house implementation of the Metropolis-Hastings (MH) sampling algorithm is used for posterior inference, available from the AmiryousefiLab/ENDS Github. Knots are chosen at the unique doses and the parameter *λ* is chosen as the value with minimum squared error over a grid of values including the mean variance estimate of the samples at each knot. We also chose a uniform Dirichlet distribution as prior for *a*, this is *a* ∼ *Dir*(1, ‥, 1), slightly different model choice from Roerink *et al*. (2018) where a stick breaking process is used to generate the weights of the Dirichlet distribution, note that a stick breaking process would force the weights to be decreasing in magnitude with probability one, we think this assumption is too restrictive, so our approach assumes a uniform Dirichlet prior which allows the weights to be any magnitude as long as they are non negative and sum up to one.

Once the samples of the posterior distribution are obtained from the chains of MH algorithm, we drop out the first half of the observations by default. Once we have the estimate of the parameters *Ĉ, â*, 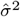 as the mean values, the posterior curve is obtained as

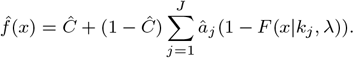

The IC_50_ value is calculated from this function as

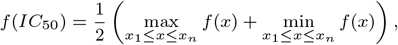

and for any *p* ∈ (0, 1) we have *IC*_*p*_ obtained as the value such that

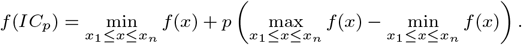

Note that the Bayesian approach would allow us to obtain for each sample of parameters an *IC* and then have a distribution for it, from which we could take the mean, but to keep the congruency with other models, we opt to obtain the IC_50_ as the cross point of the fit that would be marked over the curve in the plot.

The MH algorithm uses as priors normal distributions with parameters manually found such that the posterior sampling rate was above 30% for all parameters. Note that we have to use a transformation of the parameters such that they are in the correct range, for *σ*^2^ we used an exponential function, for *C* a logistic transformation and for *a* a softmax transformation. The change of variables rule was used to update the priors such that the MH worked correctly, this was done by adding the absolute value of the determinant of the Jacobian matrix. Lastly we chose a chain length such that 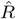 statistic is close to one for each parameter, this indicates that several chains are converging to the same values, so the chains are long enough.

#### 1.1.4 Parametric Logistic (*pL*)

This model is the logistic function adjusted to the data with four parameters found by least square estimation. The model assumes a fixed “S” shape decreasing curve for the responses. This is the usual four parameter logistic fit to the data to minimize a squared error loss function, the model fit is the logistic function given by

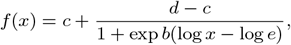

where *c* is the asymptotic minimum value, *d* is the asymptotic maximum value and *e* is the IC_50_. Note that we are using log *x* in the formula since the fit is done over the logarithm of the doses. The parameters are estimated by minimizing the squared error function ∥*y* − *f* (*x*) ∥^2^. There is not an analytical solution to this problem, different numerical minimization algorithms exist for estimating the parameters, our implementation uses the R package `drc`, which internally uses the base R optimizer `optim`. Note that the function is monotonic decreasing unless we have a degenerate fit, in which the adjusted function is a linear fit with non negative slope. Since the function is monotonic and continuous then for any value in the interval *p* ∈ (0, 1) the *IC*_*p*_ is the minimum value for which *f* (*IC*_*p*_) = *c* + *p*(*d* − *c*).

### 1.2 Web Application Options

#### 1.2.1 Input

This section shows how to start using the ENDS to fit parametric and nonparametric drug dose response curves to data. The application accepts as input a .*csv* file with the format as in Tab. 1 `Workshop` tab and provides different drug-patient characteristics for plotting (Roerink *et al*., 2018). Once the drug-response data is uploaded or the drug-patient characteristics are selected, click the `Plot` button to compile the graph.

**Table 1.**
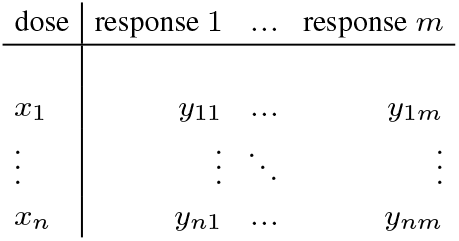
Data input accepted by web application

#### 1.2.2 Output

The plots generated can be downloaded in the `Download` tab with the `Download Plot` button. The name can be specified or the default name is the “ENDS”. The dots per inch are applicable only to *png* or *jpeg* downloads. The parameters estimated by each of the models and the statistics derived from them can be downloaded in the `Download` tab with `Download Fit` button. The post-processed data can be downloaded in the `Download` tab with `Download Data` button.

#### 1.2.3 Plot Options

The plot provides the following selective options:

##### Models

- **Nonparametric Spline (*npS*):** Simple linear spline connecting the mean responses.
- **Nonparametric Monotonic (*npM*):** Is the nonparametric spline with a non-increasing constraint added, by taking recursively the means of the previously accumulated responses until it is non-positive.
- **Nonparametric Bayesian (*npB*):** Specifies a Bayesian hierarchical model which fits a mixture of normal cumulative distribution functions centered at the doses as the basis functions.
- **Parametric Logistic (*pL*):** The usual four parameter logistic model fit by weighted least squares to the data.

##### Indices for npS

- **Point Samples:** If dataset consists of multiple samples, these are shown in the plot.
- **Min-max bands (MMB):** Linear connection of the maximum values for each dose. Same for minimum values. This could be paralleled with the confidence interval in nonparametric setting.
- **Empirical Viability Band (EVB):** It is generated by connecting the sides of the step function generated by the adjacent mean lines, which are the horizontal line that at the mean of each consequent pair of responses. It is used to visualize the variability of the samples within this bands.
- **Drug Span Gradient (DSG):** It is the linear regression fit over the mean responses, the gradient angle *θ* is given in the plot.
- **Absolute Efficiency Degree (AED):** For dose *x*_*i*+1_ is the angle of the slope of the spline from dose *x*_*i*_ to *x*_*i*+1_.
- **Relative Efficiency Degree (ARD):** Is AED−*θ*, where *θ* is the angle of the DSG. If the angle is in (0, 90) say the dose *x*_*i*_ is **Negative Relative Dose (NAD)** and otherwise it is a **Positive Relative Dose (PRD)**.
- **Maximum Effective Dose (MED):** Is the dose that compared to its previous dose, exhibit the most descent in terms of the slope (measured in degrees) of the respective section of the spline function of the fit. This is, MED is the next dose for which the minimum AED is observed.
- **Concave Ratio (CR):** Defined as *CR* = #*PRD/*#*NRD*.

##### Data processing

- **Median/Mean:** Mean or the median of the responses at that dose, by default it is mean.
- **Outliers Kept:** If not selected then samples outside of 2 standard deviations from mean are removed from data.
- **Viability over 100:** If not selected then values above 100 are replaced with 100.
- **Dose dependent AUC:** If not selected then AUC is calculated with a sequence of integers from 1 to the number of samples as the *x*-values, else with the doses as the *x*-values.
- **Spectral choice of IC:** By default set on fifty, this option allows a free choice of the IC value between zero to one hundred.

### 1.3 Comparative Analysis

For all the data found in Roerink *et al*. (2018), we have fitted each of the four models npS, npM, npB and pL to each combination of drug, patient, treatment and sample. For npB we followed the aforementioned procedure of fixing the knots of the function at the doses and choosing the variance parameter of the Normal CDF from a grid of values including the maximum likelihood estimate, as the one which minimized the squared error, and then computed the posterior parameters with the Metropolis-Hastings algorithm for 10,000 iterations. We found the IC_50_, MSE and AUC for each of the models which are shown in the Figure 1.3. Note that we have included a box plot within the violin plots to showcase where the median, first and fourth quantiles are, and outliers are marked with strong dots. Outliers are those that reside outside the first quantile minus 1.5 times the interquantile range, or the third quantile plus 1.5 the interquantile range.

Using a Shapiro test we reject the hypothesis of normality, so we use the nonparametric Wilcoxon test, to compare whether the mean of models are statistically different from each other. The results of the tests are shown in the Tab. 2, 2, and 4. In summary the IC_50_ of npS and npM are slightly smaller than the pL. The mean squared error of npS is the smallest as the mean is the statistic that minimizes the mean squared error, and so its MSE is different statistically from other models, the npB has a significantly bigger MSE than the pL. The AUC are similar in all the models.

**Table 2.** Wilcoxon test *p*-values for the IC_50_

| $IC_{50}$ | pL | npS | npM | npB |
| --- | --- | --- | --- | --- |
| pL | 1 | 0.03* | 0.04* | 0.42 |
| npS |  | 1 | 0.77 | 0** |
| npM |  |  | 1 | 0.01** |
| npB |  |  |  | 1 |

**Table 3.**
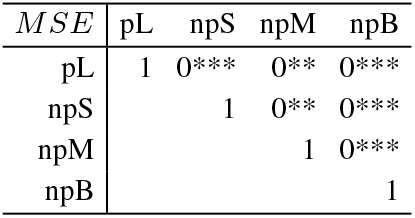
Wilcoxon test *p*-values for the MSE

| MSE | pL | npS | npM | npB |
| --- | --- | --- | --- | --- |
| pL | 1 | 0*** | 0** | 0*** |
| npS |  | 1 | 0** | 0*** |
| npM |  |  | 1 | 0*** |
| npB |  |  |  | 1 |

**Table 4.**
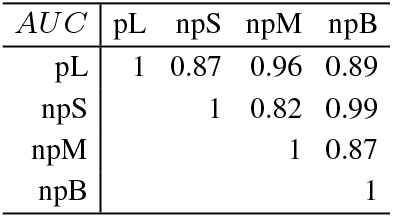
Wilcoxon test *p*-values for the AUC

| AUC | pL | npS | npM | npB |
| --- | --- | --- | --- | --- |
| pL | 1 | 0.87 | 0.96 | 0.89 |
| npS |  | 1 | 0.82 | 0.99 |
| npM |  |  | 1 | 0.87 |
| npB |  |  |  | 1 |

**Fig. 2.**
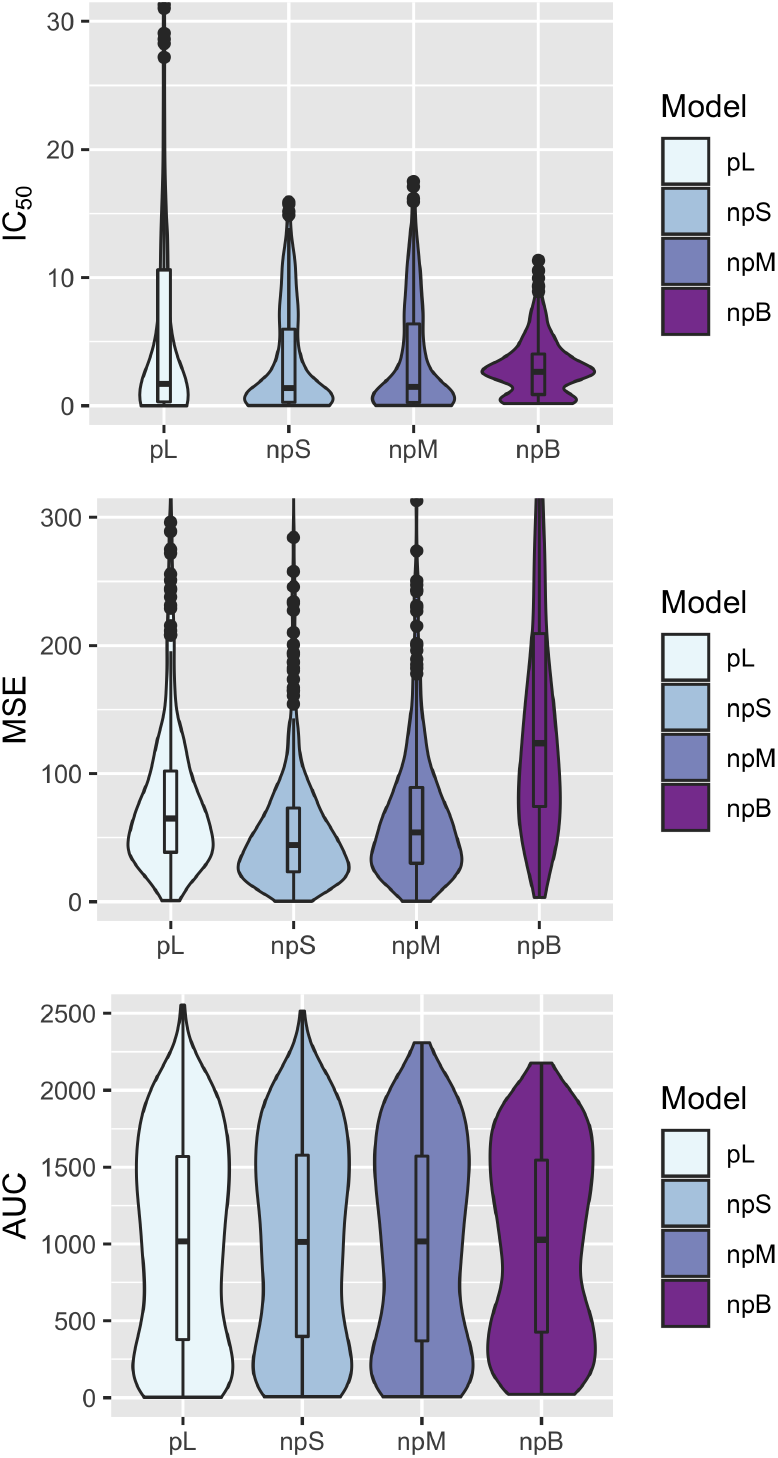
Violin plots of the IC_50_, MSE and AUC of the pL, npS, npM and npB models.

